# Separating the effects of brown-rot and white-rot fungi on crystalline cellulose using Raman spectroscopy

**DOI:** 10.1101/2025.10.06.680762

**Authors:** Ashish Ahlawat, Anders Tunlid, Dimitrios Floudas

## Abstract

Mushroom-forming wood-decay fungi are broadly categorized into white and brown rot. White-rot fungi decompose recalcitrant crystalline cellulose using a large array of hydrolytic and oxidative enzymes. Brown-rot fungi lack many of these enzymes but decompose cellulose via Fenton-generated hydroxyl radicals. To better understand these mechanisms, we developed a Raman spectroscopy-based method to study cellulose decomposition by two white-rot fungi (*Bjerkandera adusta* and *Trametes versicolor*), a brown-rot fungus (*Fomitopsis pinicola*), and a *Stereaceae* species of uncertain decay type. Raman spectra of fungi-decomposed cellulose were highly complex, reflecting physical and chemical cellulose modifications and fungal compounds like pigments. To extract signals only from decomposed cellulose and reduce data dimensionality, reference libraries were generated using chemicals that reduce crystallinity (NaOH) or oxidize cellulose (TEMPO). Chemical libraries reduced data complexity and facilitated extraction of cellulose decomposition signals, distinguishing white-rot from brown-rot effects. Wavenumbers related to oxidation better contributed to the separation of the two decomposition types. The reduced datasets also matched the decomposition characteristics of the uncertain decay fungus to those of the brown-rot fungus. The methodology developed here could be used to further characterize plant cell wall biopolymer decomposition in single fungus-single substrate setups and complex soil samples.

**Importance:** The degrading activity of saprotrophic fungi plays a crucial role in organic matter decomposition in terrestrial ecosystems, influencing nutrient cycling from plant material. Recently, there has been increasing interest in utilizing spectroscopic techniques for studying organic matter decomposition, as these methods are non-destructive. While Raman spectroscopy has been employed to identify and differentiate chemical compounds, its application to biological samples has been limited due to the complexity of spectral signals, which are challenging to interpret. In this study, we introduce a novel approach to reducing Raman spectral data to elucidate the mechanisms underlying fungal degradation of cellulose. This is achieved by utilizing reduced datasets derived from the spectral analysis of chemically modified cellulose. The dataset can be further expanded to include additional chemical treatments and fungal species, potentially revealing differences in cellulose degradation among various saprotrophs. Moreover, this approach can be adapted for use with other substrates or chemical processes and could be enhanced by integrating omics techniques.

## 1. Introduction

Cellulose, the most abundant organic compound on Earth [1], is a structural component of plant cell walls and contributes significantly to the rigidity and durability of wood. Cellulose is a linear polysaccharide composed of glucose moieties linked by β-(1→4) glycosidic bonds [2]. These linear chains assemble into microfibrils through non-covalent interactions, primarily hydrogen bonds, and further organize into fibrils in a highly ordered, hierarchical arrangement [2, 3]. This structure comprises regions of crystalline cellulose, where cellulose chains are tightly aligned, interspersed with less-ordered, amorphous regions which arise from disruptions in hydrogen bonding caused by physical or chemical modifications [4–6]. In wood, crystalline cellulose is particularly abundant, accounting for 40-60% of the dry weight of wood [7, 8]. In addition to wood, cellulose is an important carbon source in soils comprising 20-30% of the plant litter biomass [9], and contributing to 20-40 % of the generated soil glucose [10]. Additionally, soil cellulose acts as a soil aggregation agent and therefore, its persistence in soil can affect soil carbon stability [11].

The complex higher-order structure of cellulose, particularly the tightly packed arrangement of cellulose chains in the crystalline regions, makes it resistant to decay by depolymerizing enzymes [3]. Nevertheless, microbial decomposers have developed mechanisms to decompose cellulose including its crystalline regions [12]. Such microbes include wood decayers, which are mostly represented by filamentous fungi, but also many leaf litter and soil-inhabiting microbes, which readily utilize cellulose [9, 13]. Among filamentous fungi, a group of particular interest is mushroom forming wood decay species, because they cause large parts of wood and subsequently cellulose decomposition. These fungi are categorized into white rot and brown rot based on their wood decay strategies [14]. White-rot fungi can break down all major wood components—cellulose, hemicellulose, and lignin—through a suite of hydrolytic (cellobiohydrolases, endoglucanases) and oxidative (lytic polysaccharide monooxygenases (LPMOs), cellobiose dehydrogenases) enzymes [15–17]. These enzymes work in coordination to depolymerize crystalline cellulose affecting only surface crystallinity [18, 19] having little effect on the ratio between the fibril diameter and fiber crystallinity [20].

In contrast, brown-rot fungi typically lack the enzymes necessary for crystalline cellulose decomposition [21–23], though they can still extensively decompose cellulose, leaving a brittle lignin residue [24, 25]. Brown-rot fungi have evolved from white-rot ancestors [26] and during these transitions they lost many genes involved in plant cell wall decay [21–23, 26, 27] particularly those related to the depolymerization of crystalline cellulose, while maintaining gene coding for endoglucanases which act on amorphous cellulose [28].

The lack of cellulolytic activity [29, 30] and the limited capacity of brown-rot fungi to degrade lignin [31–35] has led to the suggestion of an alternative mechanism that allows full depolymerization of carbohydrates without substantial lignin removal. Accordingly, brown-rot fungi secrete small secondary metabolites [36–38] which can diffuse into the wood matrix, reduce iron, and potentially generate hydrogen peroxide [36, 39–41]. The reaction of iron and hydrogen peroxide generates short-lived hydroxyl (HO•) radicals through Fenton chemistry, which result in oxidation of wood polymers [42] including cellulose [43, 44]. Early studies suggested that brown-rot fungi rapidly decrease the degree of polymerization of cellulose by acting on the amorphous regions between the crystallites [45–47]. More recent studies have shown that brown-rot fungi also degrade the crystalline regions of cellulose - a distinct feature that contrasts with white-rot fungi, which does not reduce cellulose crystallinity to the same extent [28, 48–50]. The mechanism underlying the loss of crystallinity during brown rot decay is not clear considering that brown-rot fungi have lost many of the enzymes involved in crystalline cellulose degradation that are present in white-rot fungi [23, 51]. It has been proposed that the loss of cellulose crystallinity caused by brown-rot fungi is due to non-enzymatic processes involving the action of HO• radicals and metal-chelators [49, 50].

Raman spectroscopy has been widely used to characterize the structure of both intact and modified cellulose [52–54]. Most studies have focused on physical and chemical modifications, particularly the loss of crystallinity, but none have specifically examined cellulose oxidation using Raman spectroscopy [54–57]. Moreover, only one study has examined changes in the structure of crystalline cellulose caused by fungi, and the study focused only on a narrow region of the Raman spectrum of cellulose [28]. Analyzing cellulose modifications using Raman spectroscopy is complex, since several wavenumbers are simultaneously affected during cellulose treatment [52, 54, 56, 57]. The orientation of cellulose fibrils relative to the Raman laser affects the intensity of peaks, which can affect the interpretation of chemical and structural changes [55, 58]. The introduction of fungi adds further challenges, due to interfering signals from autofluorescence, pigments, and enzymes [59–61]. Autofluorescence arises from organic compounds emitting light as fluorescence after specific wavelength absorption [60], while fungal pigments and enzymes can produce intense Raman peaks [61].

In this study, we used Raman spectroscopy to detect changes in the chemical and physical structure of cellulose caused by two white-rot fungi (*Bjerkandera adusta* and *Trametes versicolor*), a brown-rot fungus (*Fomitopsis pinicola*), and a member of Stereaceae (Russulales) with an uncertain decay type. In contrast to an earlier study [28] that also examined fungal degradation of cellulose using Raman spectroscopy, the whole Raman spectrum of cellulose was analyzed. To disentangle spectral signatures related to loss of crystallinity from those related to oxidation as well as to remove interfering signal from biological components (e.g. hyphae, pigments, chitin etc.) we developed a data reduction strategy using reference libraries from well-characterized chemical treatments, such as sodium hydroxide (to reduce crystallinity) [62] and TEMPO (to oxidize cellulose) [63, 64]. This approach allowed us to identify key wavenumbers associated with loss of crystallinity and oxidation, and separate white-rot from brown-rot fungi, while minimizing the interference from fungi. The method provides a framework for distinguishing the effects of different fungi on cellulose and offers a basis for their classification based on their decay mechanism.

## 2. Materials and methods

### 2.1. Cellulose pellets

Cotton cellulose was purchased from Sigma Aldrich (Product no: BR28205, Lot.: 20115161). The relative crystallinity of cotton cellulose is 85 % [65]. Cellulose pellets of 150 mg with diameter of 13 mm were made using a Specac Atlas^TM^ Manual Hydraulic Press (Specac Ltd., Kent, England) applying a pressure of 10 tonnes for 1 min. The pellets were autoclaved and remained in a sealed jar until they were used for the experiments.

### 2.2. Chemical treatments of cotton pellets

Two chemical treatments were performed on cellulose. The first, aimed at inducing amorphogenesis (reducing cellulose crystallinity), used NaOH [62, 66] under conditions from Jiao and Xiong [62]. A sterile cotton pellet was placed in a small petri dish (35 x 15 mm), and 4 mL of 4M NaOH solution was added. A sterile cotton pellet was placed in a small petri dish (35 × 15 mm) with 4 mL of 4 M NaOH. The reaction occurred at room temperature and was stopped after 1 and 4 hours by removing the NaOH and washing gently with 5 × 5 mL deionized water. The pellet was dried at 50 °C for 24 hours. A small piece was transferred to a microscope slide, covered with a coverslip, and placed under the Raman spectrometer for spectra collection.

The second treatment oxidized cellulose using TEMPO (2,2,6,6-tetramethylpiperidin-1-yl)oxyl) (Sigma Aldrich) [63, 64], following Isogai et al. [63]. A sterile cotton pellet was placed in a similar petri dish with 0.0125 g each of TEMPO and sodium bromide (NaBr), and 3 mL of 9% (w/v) sodium hypochlorite (NaClO, pH 11–12). Samples were incubated at room temperature for 1, 4, or 24 hours, then washed with deionized water, dried, and a small piece was transferred to the Raman spectrometer as described for the NaOH treatment.

### 2.3. Fungal strains

Four fungal strains were used*: Bjerkandera adusta* HHB-12826 sp. (BA, white-rot fungus), *Trametes versicolor* FD-591 (TV, white-rot fungus), *Fomitopsis pinicola* FD-585 (FP, brown-rot fungus), and a member of Stereaceae (FD-574SV, Russulales, strain isolated from a corticioid fruitbody collected on well-decayed wood in Tjörnarp, Sweden) with uncertain type of decay (UWD). Fungal cultures were maintained on malt extract agar medium with trace metal solution and vitamin solution (see supplementary section A).

### 2.4. PCR, sequencing, and phylogenetic analysis

Three different DNA samples were extracted from freshly grown mycelium of the unidentified strain FD-574SV using the XNAT – REDExtract-N-Amp™ Tissue PCR Kit (Sigma-Aldrich), followed by a purification step using the ExoSAP-IT™ PCR Product Cleanup Reagent (Thermo Fisher Scientific, Waltham, MA USA). The ITS sequence was generated using the ITS1F/ITS4 primers [67, 68] using the following PCR protocol: (94°C 3 min Hot Start), 94°C 1 min, 55°C 1 min, 72°C 1.5 min, 35 cycles. The sequencing reactions and sequencing were done by GeneWiz Europe (Azenta, Griesheim, Germany). The generated ITS sequences were used for blast searches in GenBank recovering 250 sequences in total. After a preliminary alignment followed by a neighbor-joining (NJ) tree [69] the dataset was reduced by removing identical ITS sequences. The final dataset contained 144 sequences, the FD-574SV ITS sequence, and an ITS sequence of *Trametes versicolor* (Polyporales) that was used as outgroup. The sequences were aligned using FFT-NS-i (Slow; iterative refinement method) in MAFFT [70]. The alignment was manually examined and edited in Jalview [71] to remove poorly aligned regions. The final dataset contained 655 characters in total. NJ analysis was performed in MEGA-X [72] with 300 bootstrap runs and default settings.

### 2.4. Fungal treatments of crystalline cellulose

A sterile cotton pellet was placed at the center of agar petri dishes (90 x 16 mm) containing Highley agar medium (pH 4.8) [28], (see supplementary section A), 10 mL of a vitamin stock solution and 15 g agar. A mycelial plug (c. 5 mm in diameter) was cut from the margin of a mycelium grown on the malt extract agar medium. The plug was placed near the cotton pellet and the petri dish was incubated at 25 °C for 4 weeks in darkness. At this time, the cotton pellet was fully colonized by the fungal mycelium. The cellulose pellet with adhering mycelium was removed, dried at 50 °C for 24 hours and a small piece was transferred to a microscope slide, covered with a coverslip, and placed under the Raman spectrometer for measurements.

### 2.5. Raman spectroscopy

A Horiba LabRAM HR Evolution Raman spectrometer (HORIBA, Longjumeau, France) was used equipped with a green laser (λ = 532 nm) with 600gr/mm grating. Wavenumber calibration was performed using a silicon sample.

Acquisition time (duration of laser exposure and signal collection) was set to 7 seconds, and accumulation (spectral repetitions) to 7. An optical microscope attached to the spectrometer was used to focus on cellulose fibers, and spectra were recorded from 12 randomly selected points, considered as replicates. Spectra were measured either continuously (250-3600 cm^-1^) or in six different spectral regions (60-250 cm^-1^, 250-600 cm^-1^, 820-945 cm^-1^, 945-1225 cm^-1^, 1225-1495 cm^-1^, and 2800-3600 cm^-1^) [52, 53, 55, 57].

### 2.6. Data analysis

The workflow for analyzing the Raman data of reference libraries is outlined in Fig. 1. First, spectra measured from different regions were merged to create an extended spectrum, followed by rubberband baseline correction and vector normalization (scaling the spectrum so that maximum intensity is 1 without altering relative intensities) using Quasar (Orange Spectroscopy) [73]. For comparing the spectra, the spectral files were imported and visualized using OCTAVVS [74]. Identification of peaks was performed using the iPeak script in MATLAB [75] with the following settings: Gaussian peaks with an amplitude threshold of 0.005, slope threshold of 2.6e-05, smooth width of 2, and fit width of 2.

**Figure 1.**
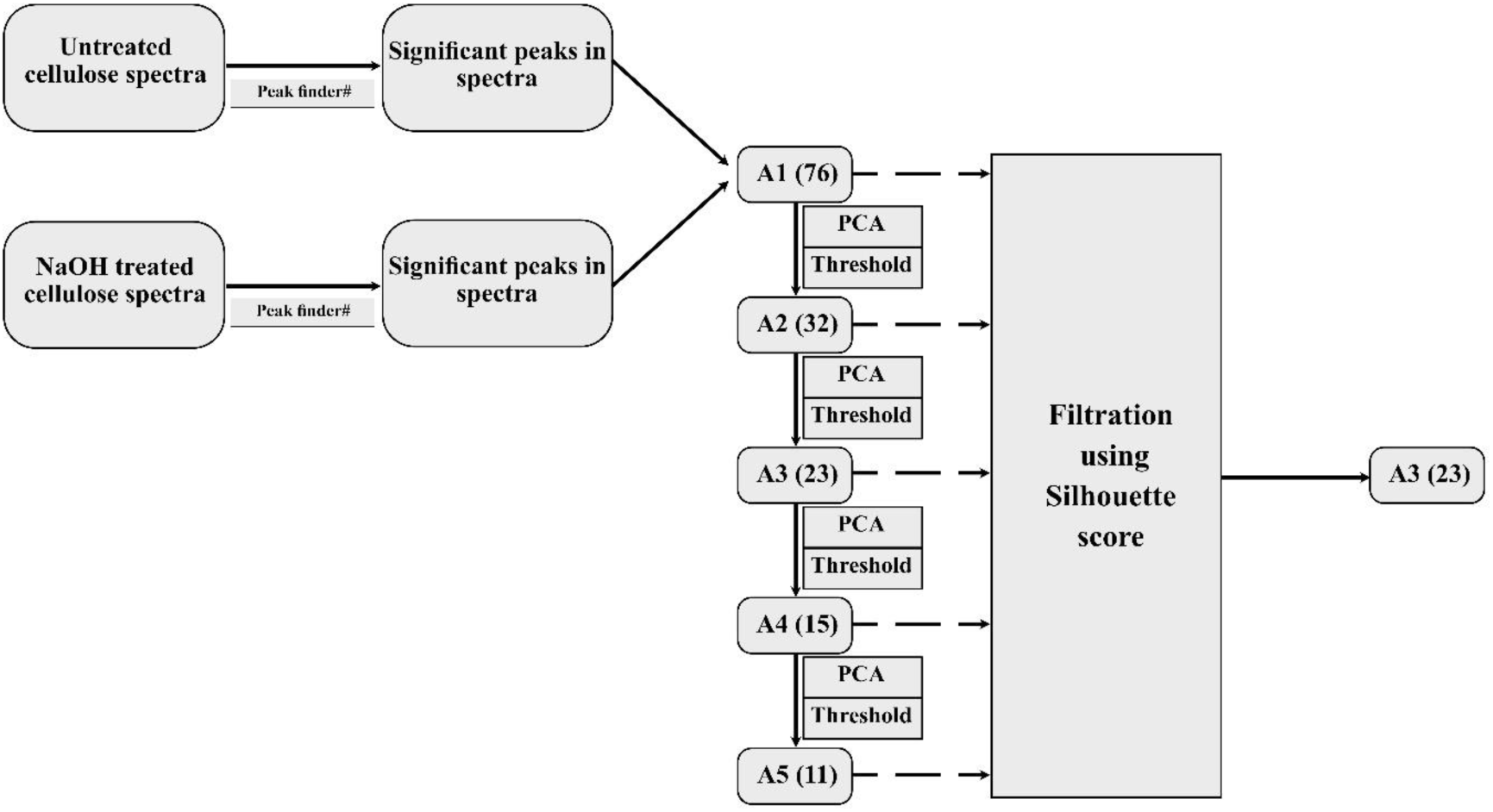
Workflow for NaOH-modified cellulose Raman data processing and reduction. Cellulose was chemically treated to induce amorphogenesis (using NaOH). Spectra were collected for both treated and untreated cellulose across six regions and preprocessed. After selecting significant peaks using Peak finder, the data was then progressively reduced using PCA loading values, resulting in reference libraries of wavenumbers associated with amorphogenesis (A1-A5). Subsequently, the A3 library was selected based on silhouette score threshold. Similar workflow was followed to generate oxidation libraries.

To extract the signals from cellulose and reduce the dimensionality of the data, two sets of reference libraries were constructed: Amorphogenesis (A) libraries containing wavenumbers that are indicative for the loss of crystallinity, and oxidation (O) libraries with wavenumbers associated with oxidation of cellulose. Data reduction was performed using Principal Component Analysis (PCA) using the R Statistical Software v4.1.2 [76] in R Studio. The data input for generating the A libraries contained the wavenumbers and intensities of peaks identified in the samples treated with NaOH and those of nontreated cellulose (Control). For the O libraries, the input data were wavenumbers and intensities from the TEMPO treated and nontreated cellulose samples. PCA was conducted, and the loading values of principal components 1 and 2 (PC1 and PC2) were used to identify wavenumbers contributing most to the separation between NaOH/TEMPO treated and untreated cellulose. To isolate wavenumbers maximizing the separation, thresholds were applied to the loading values based on the magnitude of PC1 (≥ 0.05 or ≤ -0.05) and the directions of PC1 and PC2 (tan⁻¹(PC2/PC1) between -45° and 45°). This process was repeated iteratively to generate five A (A1–A5) and five O (O1-O5) libraries containing decreasing numbers of wavenumbers. The best libraries were identified based on the Silhouette score [77], which measures the mean intra-cluster distance and the mean nearest-cluster distance for each sample. The distances were calculated using the PC1 and PC2 score values. The highest Silhouette scores were obtained for the A3 and O3 libraries, and their wavenumbers were categorized into three groups: A* (unique wavenumbers in A3), O* (unique wavenumbers in O3), and C* (common wavenumbers in both A3 and O3). The wavenumbers in these groups were then used to reduce the Raman spectral data of fungal-decomposed cellulose. The intensities of specific spectral peaks (1460/1481 cm^-1^, 376/1095 cm^-1^, 90 cm^-1^) were used to assess the crystallinity in cellulose [54, 78, 79].

### 2.7. Statistical tests

PCA was performed on fungal treated and untreated data. The scores of PC1 and PC2 were used for calculating the Euclidean distances between the samples of fungal-treated and non-treated cellulose. For any given wavenumber, its contribution to separating the treatments was evaluated by calculating the distances (i.e. sum of the Euclidean distances) between the samples when the wavenumber was present or excluded from the analysis. The decrease in distance (%) due to the removal was used as a measure of the wavelength’s contribution. This calculation was performed for all wavenumbers. Contributions were normalized to the value representing the largest distance decrease. A mixed-effects model [80] was fitted to the reduced fungal data to test species differences, followed by Bonferroni-adjusted pairwise comparisons. Results were visualized in a heatmap. To test MANOVA’s multivariate normality assumption, the Shapiro-Wilk test [81] was used to detect outliers. Differences between white-rot and brown-rot species were as-sessed using Discriminant Function Analysis (DFA) with MANOVA. The candisc package [82] illustrated the role of specific wavenumbers in distinguishing species.

## 3. Results

### 3.1. Preprocessing of Raman spectra of cellulose colonized by fungi

Cotton pellets containing highly crystalline cellulose were colonized by two white-rot fungi (*B. adusta* and *T. versicolor*), one brown-rot fungus (*F. pinicola*) and an isolate (FD-574SV) with an uncertain type of wood decay (UWD). The ITS sequence of the latter strain resulted in no close matches in GenBank (closest hit was 88%), and the closest hits belonged to the genera *Conferticium*, *Xylobolus*, and *Stereum*. In the resulting NJ tree including the closest ITS sequence hits from Genbank, FD-574SV took a basal placement in a well-supported clade containing *Xylobolus* sequences (Fig. S1). The low match of the strain with any of the genera recovered from Genbank and the lack of a fruitbody from that collection makes the genus or species assignment of the strain impossible. However, our results support the placement of the species among the corticioid members of Stereaceae [83] (*Xylobolus*, *Stereum*, *Acanthophysium*, *Aleurodiscus*, *Conferticium*).

The Raman spectra of the fungi-decomposed cellulose were initially recorded continuously from 250 to 3600 cm⁻¹. These measurements revealed strong autofluorescence signals (Fig. 2A) [59] and, in the case of the fungus with unknown decay type, distinct peaks at 995, 1130, 1521, 2256, 2638 and 3033 cm^-1^ attributed to carotenoids (Fig. 2B) [61]. To minimize interference from such compounds and enhance the signals from cellulose the spectra were measured in four regions containing characteristic cellulose peaks [52, 54, 78, 79]. One of them (820–1495 cm⁻¹) was further split into three regions to reduce strong signal from carotenoid peak at 1160 cm^-1^ (Fig. 2B). The merged spectrum included peaks corresponding to 768 wavenumbers. Even though, the signals from cellulose were enhanced in these spectra, a PCA revealed poor separation between fungal-treated cellulose and the non-treated cellulose (control) (Fig. 2C). Moreover, the variation among replicates was large, particularly for samples colonized by *F. pinicola* and *T. versicolor*.

**Figure 2.**
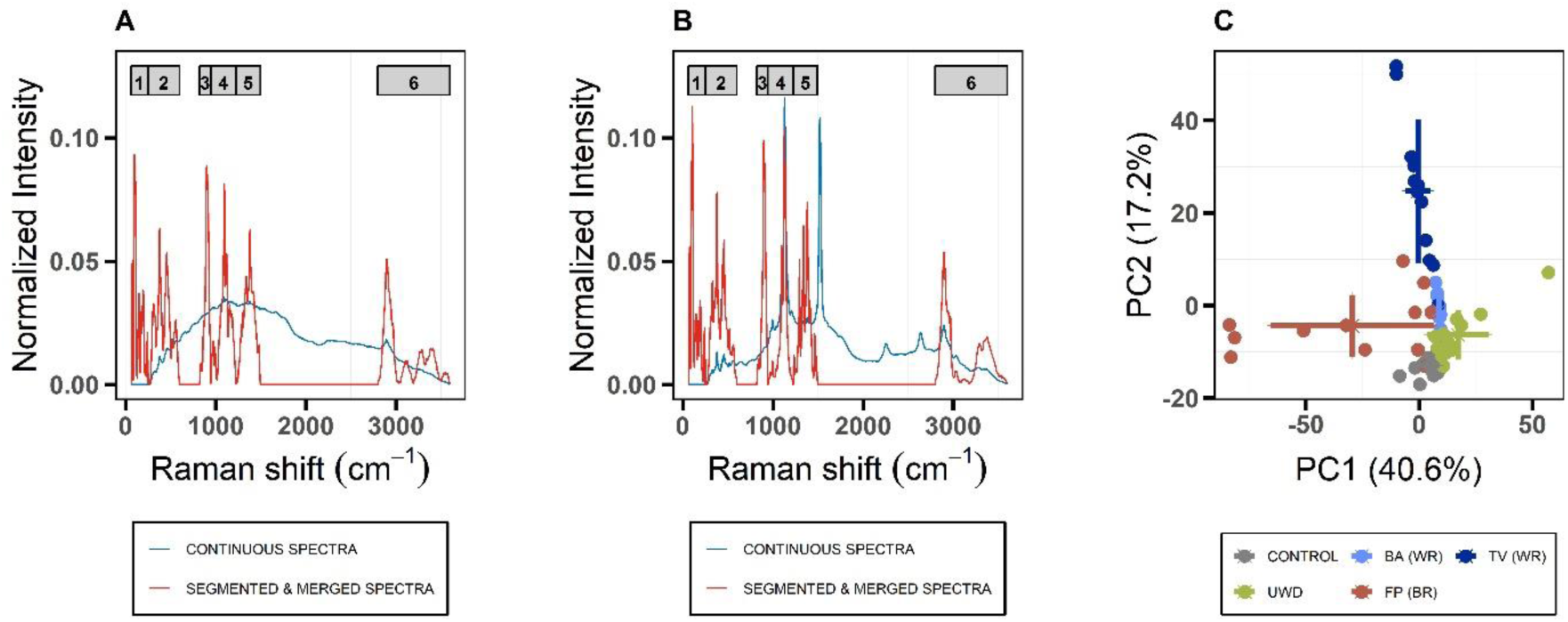
Raman spectra preprocessing. Raman spectra of cellulose colonized by wood-decay fungi were measured either continuously from 250 to 3600 cm⁻¹ (blue) or in six selected regions merged into a single spectrum (red). The six regions, indicated by grey bars, were 1) 60-250 cm⁻¹, 2) 250-600 cm⁻¹, 3) 820-945 cm⁻¹, 4) 945-1225 cm⁻¹, 5) 1225-1495 cm⁻¹, and 6) 2800-3600 cm⁻¹. (A) The average spectrum of cellulose colonized by *Trametes versicolor* (TV) highlights the need to avoid certain regions (600-850 cm⁻¹) to reduce strong background signal from fungal colonization. The comparison between the continuously measured and regionally measured, merged spectrum shows only a few discernible cellulose peaks (c.f. Fig. 4). (B) The average spectrum of cellulose colonized by strain FD-574SV that segmented measurements can facilitate avoiding high-intensity carotenoid peaks (1129 cm⁻¹ and 1520 cm⁻¹). (C) A PCA plot of merged spectra from four wood-decay fungi species shows that PC1 and PC2 components explain 57.8% of the variance. Each point represents spectral data from one sample. Each treatment has data from 12 samples (replicates). The cross for each label indicates the standard deviation across the PC1 and PC2 scores. WR, White-rot fungi; BR, Brown-rot fungi; BA, *B. adusta*; TV, *T. versicolor*; UWD, a strain of a species of unknown wood-decay type; FP, *F. pinicola*.

### 3.2. Generation of reference chemical libraries

To identify spectroscopic signals from cellulose associated with amorphogenesis and oxidation, reference libraries of wavenumbers were generated using cellulose treated with NaOH [62, 66] and TEMPO [63, 64], which are known to induce amorphogenesis and oxidation, respectively. PCA of cellulose treated with NaOH for 1 and 4 hours revealed distinct separation between NaOH-treated and non-treated cellulose, but no separation was observed between the 1-hour and 4-hour NaOH-treated samples (Fig. 3A). The lack of separation between 1-hour and 4-hours treatments suggests that most modifications caused by NaOH were completed already during the first hour of incubation, even though it should be noted that samples from 1-hour treatment are more variable than samples of the 4-hours treatment. The PCA dataset included spectroscopic signals from 768 wavenumbers (A0 library). To reduce dimensionality, a sequential data reduction strategy was applied, resulting in five amorphogenesis libraries (A1–A5) (Fig. 3). Among these, the A3 library, containing 23 wavenumbers, had the highest Silhouette score (0.52), indicating that the best separation between NaOH treated and non-treated cellulose was achieved using the wavenumbers of this library (Fig. 3B). The explained variance in the plots (Fig. 3A and C) increase considerably after data reduction.

**Figure 3.**
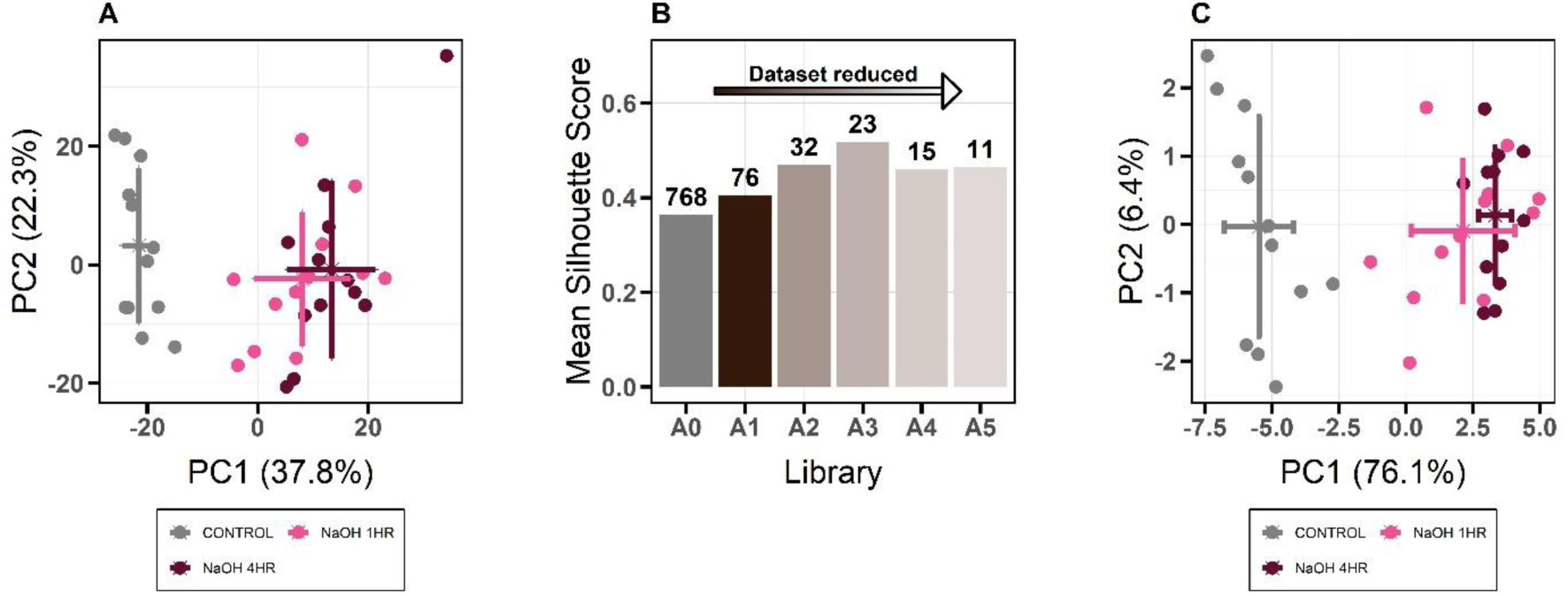
Generation of reference libraries associated with loss of crystallinity in cellulose. (A) PCA plot of Raman spectra from non-treated cellulose (control) and NaOH-treated cellulose (for 1 and 4 hours) before data reduction (A0 library, with 768 wavenumbers). PC1 and PC2 explain 60.1% of the total variation. Each point represents data from one replicate (n=12), and the cross indicates standard deviation across the PC1 and PC2 scores. (B) Silhouette scores of libraries (A1 to A5) generated by reducing the number of wavenumbers in the A0 library using a sequential analysis of PCA loading values. The number of wavenumbers in these libraries is indicated above the bars (decreasing from left to right). (C) PCA plot of spectra from non-treated and NaOH-treated cellulose using data from the 23 wavenumbers in the A3 library. PC1 and PC2 explain 82.5% of the total variance, with each point representing data from one replicate (n=12), and the cross representing standard deviation across the PC1 and PC2 scores.

A similar approach was used to identify oxidation-associated signals from cellulose treated with TEMPO (Fig. S2). The complete data set contained spectroscopic signals from 768 wavenumbers (O0 library), and the data reduction strategy generated five oxidation libraries (O1–O5). The highest Silhouette score (0.57) was obtained for the O3 library which contained 23 wavenumbers. A similar increase in the explained variance was observed in PCA plots of reduced library (Fig. S2A, S2C). There were no significant differences between cellulose treated with TEMPO for 1, 4, or 24 hours (Fig. S2A) suggesting that most cellulose modifications were already completed during the first hour of incubation.

Wavenumbers from the A3 and O3 libraries were grouped into three datasets: A* (amorphogenesis-specific), O* (oxidation-specific), and C* (common to both A3 and O3). The A* and O* datasets each contained 12 unique wavenumbers, while the C* dataset included 11 shared wavenumbers, indicating overlap between oxidation- and amorphogenesis-associated signals. In Fig. 4, the wavenumbers from these libraries are mapped onto a Raman spectrum of crystalline cellulose. Of the 35 peaks identified in the A*, O* and C* libraries, 11 have previously been identified to be modified during amorphogenesis of crystalline cellulose, and in addition, 13 peaks have been assigned in Raman spectra of cellulose (Table S1).

**Figure 4.**
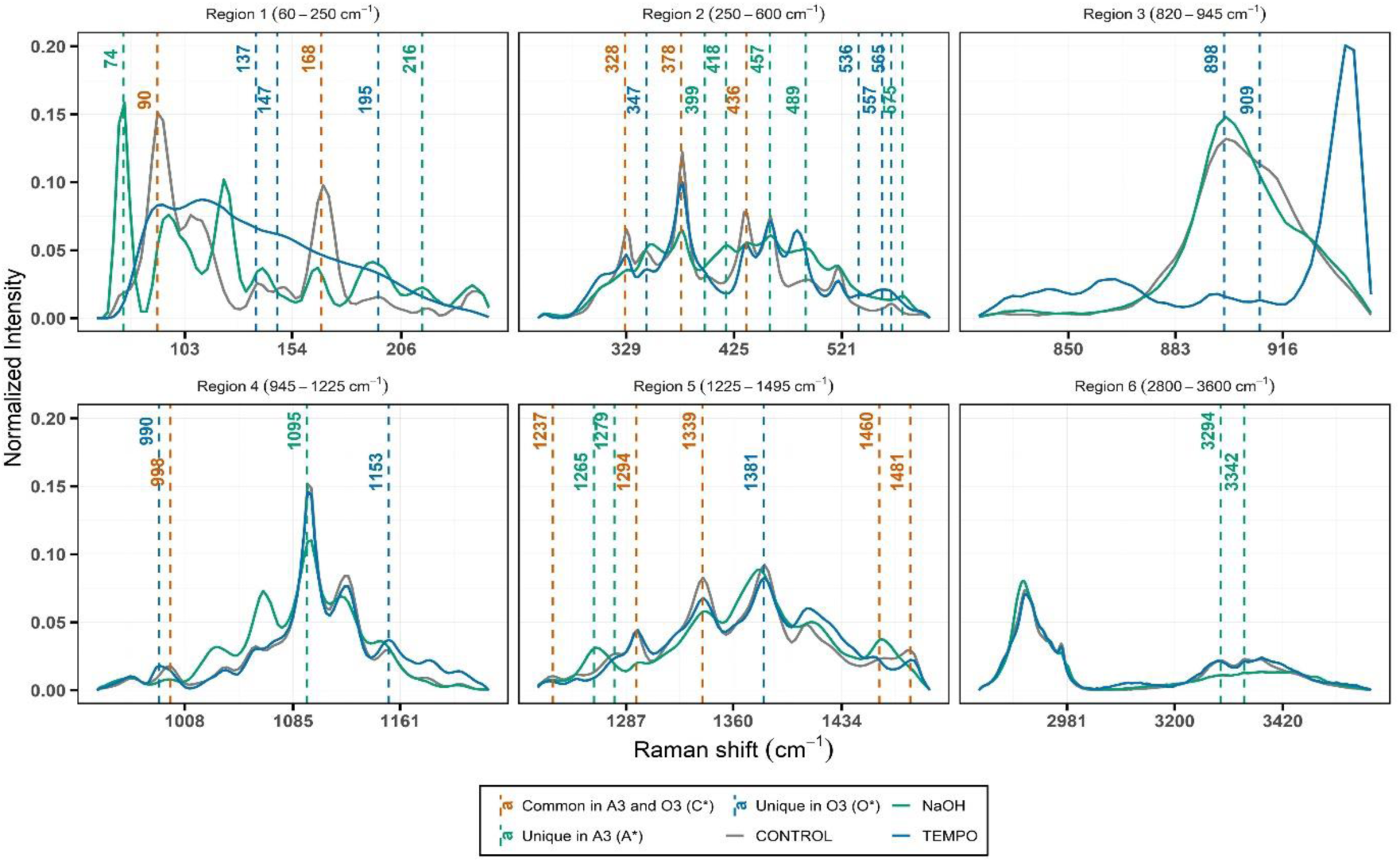
Raman spectra of cellulose associated with amorphogenesis and oxidation. The plot shows Raman spectra separated into six different regions. The average spectrum of crystalline cellulose (control, n=12) is shown in grey; the average spectrum of all NaOH-modified cellulose (n=24) is shown in green, and the average spectrum of all TEMPO-modified cellulose (n=36) is shown in blue. Wavenumbers were separated into three datasets: A*, Amorphogenesis-specific wavenumbers (unique for the A3 library, indicated in green); O*, Oxidation-specific (unique for the O3 library, indicated in blue); and C*, Common wavenumbers that are present in both A3 and O3 (orange).

### 3.3. Differentiating fungal decay types

Using the 35 wavenumbers present in the A*, O* and C* libraries, PCA successfully separated the four fungal species from untreated cellulose (Fig. 5A). PC1 and PC2 explained 51.5% of the total variance. In comparison with the PCA of the original, non-reduced dataset (c.f. Fig. 2C), the explained variance decreased, but the clustering of replicates and the separation of fungal species and untreated cellulose improved. Principal components 3 and 4 explained 20 % of the total variance, and the separation of the species was less distinct than in the plot of PC1 and PC2 (Fig. S3). Scatter plot of PC3 and PC4 from Raman spectra of fungal treated and non-treated cellulose are shown in Fig. S4.

**Figure 5.**
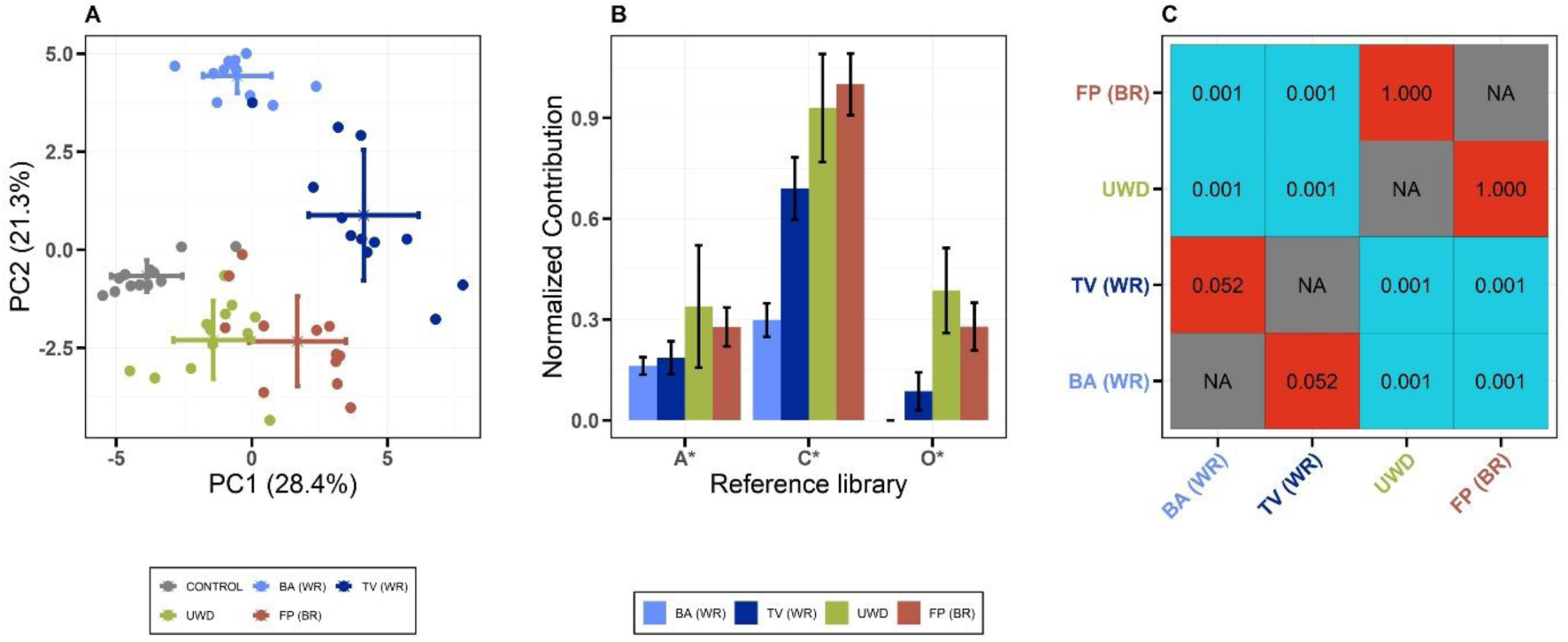
Raman spectroscopy of fungal-decomposed cellulose. (A) PCA plot of reduced spectral data from Raman spectra colonized by four different fungal species and untreated cellulose. In total, 35 wavenumbers in the A*, O*, and C* libraries were analyzed. Each point represents data from one sample, with each treatment carried out with 12 replicates. The cross indicates the standard deviation across PC1 and PC2 scores. (B) Contribution of wavenumbers from the A*, O*, and C* libraries for separating spectra from the four fungal species. Euclidean distances between fungal-treated and non-treated cellulose samples were calculated using the scores from PC1 and PC2, and the contribution of a given wavelength to the separation of the samples was estimated by recording the change in distance before and after removing the wavelength from the distance analysis. Values are normalized to the value representing the largest decrease in distance. Error bars represent standard deviation across replicates (n=12). (C) Pairwise comparison of spectral data from fungi-decomposed cellulose. In the heatmap, red indicates no significant differences (p-value ≥ 0.05), while blue indicates significant differences (p-value < 0.05). WR, White-rot fungi; BR, Brown-rot fungi; BA, *B. adusta*; TV, *T. versicolor*; UWD, Species of unknown wood decay type species; FP, *F. pinicola*.

Analyzing the contributions of the wavenumbers to the clustering revealed that the oxidation dataset (O*) clustered cellulose modified by brown-rot fungus separated from the white-rot fungi. The cellulose modification caused by the UWD species clustered closely with the brown-rot fungus (Fig. 5A). In contrast, the wavenumbers in the A* dataset showed limited ability to distinguish the brown-rot fungus from the white-rot fungi, while the common dataset (C*) effectively separated the white-rot fungus *B. adusta* from the other species (Fig. 5B). Further analysis using a mixed-effects model on the reduced fungal data confirmed significant separation between the two white-rot fungi and the brown-rot fungus (Fig. 5C). Again, the cellulose modification characteristics of the UWD species were also significantly separated from the white-rot fungi and exhibited greater similarity to the brown-rot fungus. Discriminant function analysis revealed that the two white-rot fungi were similar to each other and were distinctly separated from the brown-rot fungus (Fig. 6). However, in this analysis the UWD strain showed large differences from the white-rot fungi, but also small differences from the brown-rot fungus.

**Figure 6.**
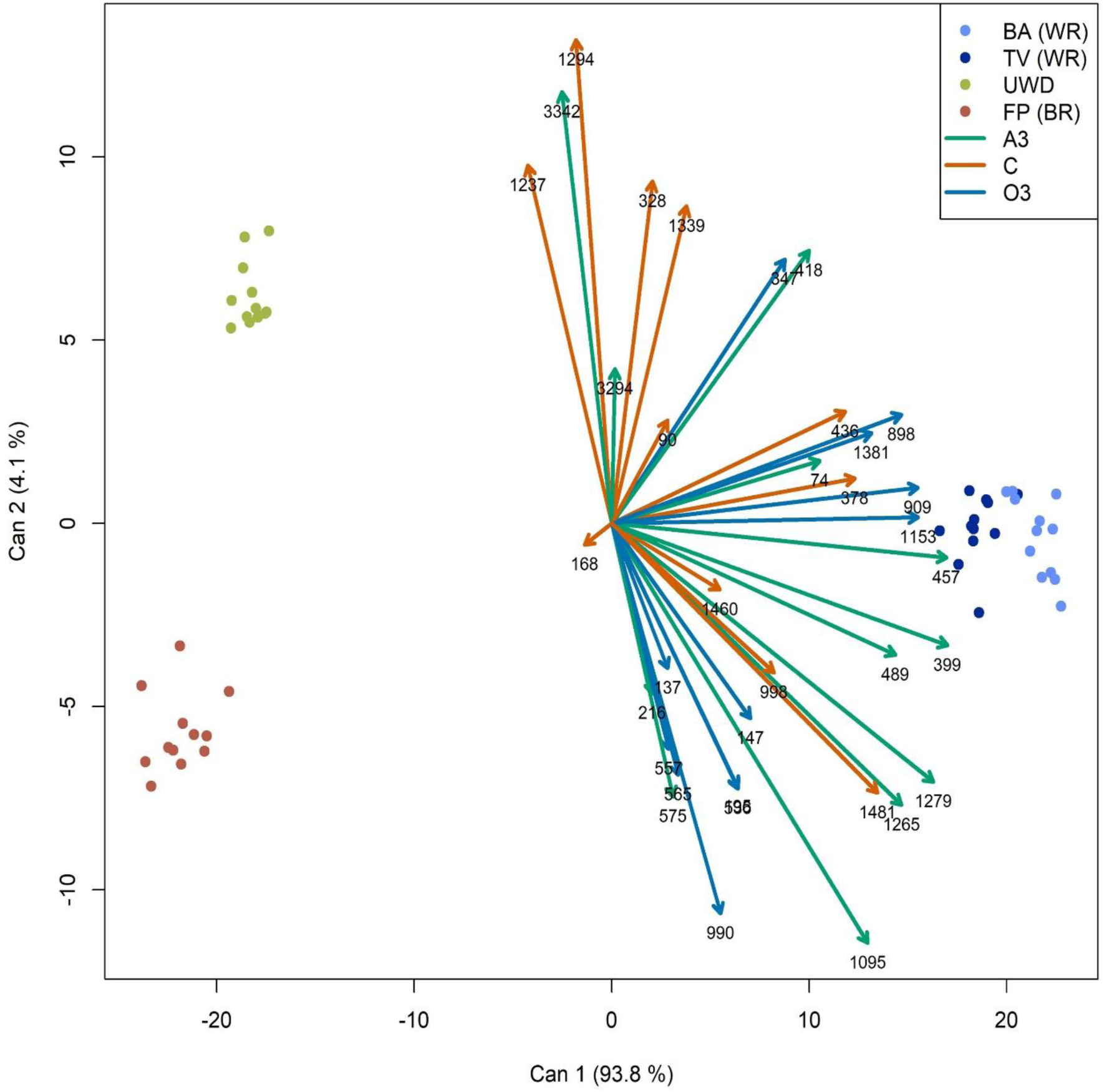
Canonical discriminant plot after performing MANOVA on reduced spectral data of fungi decomposed cellulose. Reduced spectral data of cellulose decomposed by BA & TV (WR) separated from reduced spectral data from FP (BR) and UWD. UWD is also separated from FP (BR). The wavenumbers contributing to the separation of the reduced fungi-decomposed cellulose data are shown. WR: White-rot fungi, BR: Brown-rot fungi, BA: *Bjerkandera adusta*, TV: *Trametes versicolor*, UWD: species of unknown wood decay type, FP: *Fomitopsis pinicola*.

We also assessed the effects of the fungi on the cellulose using peak intensity and peak intensity ratios from the Raman spectra peaks that have previously been identified to be associated with the degree of crystallinity and which were present in the A* and C* dataset including the 1460/1480 cm⁻¹ and 376/1095 cm⁻¹ ratios and the 90 cm⁻¹ peak (Fig. S5) [54, 56, 57]. Our results show that the intensity of the 90 cm⁻¹ peak cannot distinguish brown-rot from white-rot fungi albeit the decrease in intensity is more pronounced in the NaOH treatment rather than the TEMPO treatment. In contrast, the two peak ratios are more informative separating TEMPO from NaOH treatments suggesting that TEMPO has a much smaller effect on the intensity of these peaks and therefore a much smaller contribution to loss of crystallinity. The brown-rot fungi related ratios show that indeed brown-rot fungi tend to produce ratios that are more similar to the NaOH treatment especially for the 376/1095 cm⁻¹ pair, while white-rot fungi and especially *Bjerkandera adusta* (BA) either do not change the ratios in comparison to the control or the change is to the opposite direction from that of NaOH and brown-rot fungi.

## 4. Discussion

Wood decay mushroom forming fungi are separated into white rot and brown rot based on the chemical composition and the appearance of the decayed wood [14, 25]. The differences in crystalline cellulose decomposition by the two groups of fungi has been less clear particularly because of our limited understanding of the mechanism employed by brown-rot fungi [42]. We developed a novel non-destructive methodology to assess crystalline cellulose decomposition by fungi using Raman spectroscopy. Raman spectroscopy has been widely used to characterize cellulose [53, 55]. Moreover, so far only one study has applied Raman spectroscopy to investigate fungal decomposition of cellulose, which was focused on a small region of the cellulose Raman spectral signature [28]. This study has expanded previous work regarding cellulose decomposition by fungi by utilizing the whole Raman spectrum of cellulose and taking into consideration the effect of oxidation. Using an iterative PCA-based method based on the analyses of chemical libraries related to amorphogenesis and oxidation, we have successfully reduced the number of wavelengths needed to separate the effect of white-rot from brown-rot fungi on crystalline cellulose. The fact that the majority of the wavenumbers in the reduced dataset (31 out of 35) have been identified in earlier studies examining changes in the structure of cellulose (Table S1) [53, 55, 79] supports that the developed procedure for extracting informative Raman signals from fungal-decomposed cellulose is feasible. This is the first study examining the effects of oxidation on cellulose using Raman spectroscopy. Our analysis identified 12 unique wavenumbers that can be associated with TEMPO-oxidation. In addition, we identified 6 wavenumbers that previously have not been associated with amorphogenesis of crystalline cellulose (Table S1).

Enzymatic decomposition of cellulose as performed by white-rot fungi has not been associated with large losses in crystallinity [49, 84]. By contrast, studies have shown that cellulose that is being decomposed by brown-rot fungi gradually loses its crystallinity [51, 84, 85], albeit not all brown-rot fungi have the same effect on crystalline cellulose [20].

Notably, our results show that intensities of amorphogenesis specific wavenumber did not separate the two types of decays under the conditions used here. Instead, wavenumbers in the oxidation library contributed most to the separation between white-rot and brown rot fungi. In white-rot fungi, oxidation of cellulose is mainly caused by enzymes in the LPMO family [86]. LPMOs are present in very low numbers or completely absent in brown-rot fungi [23, 28], and these fungi, oxidize cellulose by HO• radicals generated in the non-enzymatic Fenton reaction between Fe(II) and H_2_O_2_ [42]. Notably, recent studies have shown that the oxidation mechanism of LPMO involves Fenton-like chemistry [87]. However, the extent and mode of actions HO• radicals generated by LPMOs and the non-enzymatic Fenton reaction differ. In the LPMO reaction, the Fenton chemistry is confined to the binding site between the enzyme and cellulose substrate, and the HO• oxidation leads to specific cleavages of glycosidic bonds and formation of a narrow set of oxidation products [88]. In contrast, HO• radicals in the non-enzymatic Fenton reaction, is generated at the sites of interaction between Fe(II) and cellulose fibrils, and oxidation products formed is diverse and large [89, 90]. TEMPO selectively oxidizes hydroxyl group at the 6-position of the glucose moieties [63], and the formation of products with new carboxyl groups are part of the modifications caused by both LPMOs and non-enzymatic Fenton reactions [88, 90]. Whether the separation of white-rot from brown-rot fungi based on the oxidation specific wavenumbers is the result of the separate action of HO• radicals generated by LPMOs and non-enzymatic Fenton reactions or due to differences in the magnitude of oxidation warrants further investigation. Further insights into these oxidation mechanisms can be obtained by generating chemical libraries of crystalline cellulose treated with LPMOs and Fenton oxidation *in-vitro*.

The result showing that the amorphogenesis specific wavenumber did not separate the two types of decays does not imply that effects on cellulose crystallinity caused by white-rot and brown-rot fungi were similar or not significant. Our previous study focused on a limited region of the Raman spectrum containing wavenumbers that have been used to estimate crystallinity in cellulose [28, 57, 79] showed that the changes caused by the brown-rot fungus *Gloeophyllum* sp. (strain OMC-1627) were like those of NaOH treatment, whereas the effect of the white-rot fungus (*Phanerochaete* sp.) was not significant [28]. A similar analysis of the data from this study suggests that *F. pinicola* decreased crystallinity, but to a lesser extent than *the Stereaceae strain* (FD-574SV), and the effects of the white-rot species were limited (Fig. S5). Applying X-Ray scattering, we have recently shown that the examined strains of *Gloeophyllum* sp. and *F. pinicola* affect the amorphogenesis and degrade crystalline cellulose using two different mechanisms [20]. The mechanism of *Gloeophyllum* sp. involves amorphogenesis of crystalline cellulose without substantial thinning of the cellulose fibers, whereas the mechanism of *F. pinicola* causes simultaneous microfibril thinning and decreased crystalline cellulose resembling enzymatic degradation of cellulose. Differences in the changes of crystallinity caused by brown-rot fungi have also been observed using X-ray diffraction and NMR spectroscopy [49, 91]. In contrast, Raman spectroscopy is largely a surface technique, which implies that changes in the crystallinity in the interior of the fibers like those we observed for brown-rot fungi using X-ray scattering [20] will not be accounted using Raman spectroscopy. In addition, loss of crystallinity on the surface of the fibers by white-rot fungi (e.g. action of LPMOs) or due to Fenton radicals by brown-rot fungi might result in a very similar signal related to loss of crystallinity resulting in a lack of separation between white-rot and brown-rot fungi. Further studies comprising a larger set of fungal species are needed to reveal whether Raman spectroscopy can be used to separate the brown-rot decay into several distinct mechanisms.

While white rot and brown rot have clear macroscopic features that make their separation in the field easy [92], it is frequently more difficult to attribute the decay to a specific fungal species, especially if multiple fruitbodies are found on the same wood sample. This problem is very frequent for corticoid fungi. The decay strategies of the latter group of fungi have been studied very little and their type of decay is either not reported at all or it is determined indirectly based on enzymatic tests or on field observations [93, 94]. This poses a challenge because some corticioid fungi might be secondary wood decomposers or frequently grow along with other wood decayers or they might use wood as a solid substrate to produce a fruitbody without causing the reported decay. The potentially undescribed species included in this study (FD-574SV) showed cellulose degradation characteristics that resemble those of the distantly related brown-rot fungus *Fomitopsis pinicola*. FD-574SV belongs in Stereaceae, a family that includes species traditionally considered to cause a white rot [93]. However, while *Stereum* and *Xylobolus* are the two most studied white-rot genera of the family, the remaining genera and their decay strategies remain obscure. Furthermore, even between *Xylobolus* and *Stereum* there are differences in the type of decay they cause, since *Xylobolus* causes a pocket rot, which is a type of white rot connected to selective delignification [94, 95]. Our results suggest that the assignment of decays especially for little-studied species requires more attention and that species with cellulose decomposition mechanisms that resemble brown-rot might exist among the corticioid species in Stereaceae. Genome sequencing has revealed that the traditional white-rot and brown-rot dichotomy rather harbors species that represent a gradient of decay potential with diverse sets of genes not only for cellulose, but also for hemicellulose and lignin decomposition [28, 96]. How such variation contributes to the chemical and structural modifications of cellulose or other plant cell wall biopolymers is not understood, but our results show that Raman spectroscopy could be used to study this functional diversity [97, 98]. Beyond the detailed characterization of plant cell decomposition by single fungi using Raman spectroscopy, our method provides a novel methodology that can be used to examine cellulose or other biopolymers degradation in complex environments such as soils or aquatic systems where multiple organisms might act simultaneously or in succession. Such experimental set ups might offer insights and more quantifiable estimates [99] on the fate of organic carbon in soil.

## 5. Acknowledgements

This work was supported by funding from the Swedish Research Council (VR 2019-04495) and the Crafoord foundation (Grant no. 20200759). We would also like to thank Martí Pla I Ferriol for the sequencing of the molecular marker used in the study.

## 6. Conflict of interest declaration

The authors declare that they have no known competing financial interests or personal relationships that could have appeared to influence the work reported in this paper.

## 7. Authors contributions

Conceptualization: Ashish Ahlawat, Dimitrios Floudas, Formal analysis: Ashish Ahlawat, Dimitrios Floudas, Funding acquisition: Dimitrios Floudas, Investigation: Ashish Ahlawat, Methodology: Ashish Ahlawat, Anders Tunlid, Dimitrios Floudas, Supervision: Dimitrios Floudas, Writing – original draft: Ashish Ahlawat, Writing – review & editing: Ashish Ahlawat, Anders Tunlid, Dimitrios Floudas

## 8. Data access statement

All data associated with the manuscript is available here: https://data.mendeley.com/datasets/9fc2n37fzp/1

## References

1. Field CB et al. Primary production of the biosphere: Integrating terrestrial and oceanic components. Science 1998;281:237–240. 10.1126/science.281.5374.237

2. Nishiyama Y, Langan P, Chanzy H. Crystal structure and hydrogen-bonding system in cellulose Iβ from synchrotron X-ray and neutron fiber diffraction. Journal of the American Chemical Society 2002;124:9074–9082. 10.1021/JA0257319

3. Horikawa Y. Structural diversity of natural cellulose and related applications using delignified wood. Journal of Wood Science 2022;68:1–9. 10.1186/S10086-022-02061-2

4. Ciolacu D, Ciolacu F, Popa VI. Amorphous cellulose - Structure and characterization. Cellulose Chemistry and Technology 2011;45:13–21.

5. Kulasinski K et al. A comparative molecular dynamics study of crystalline, paracrystalline and amorphous states of cellulose. Cellulose 2014;21:1103–1116. 10.1007/S10570-014-0213-7

6. Bregado JL et al. Amorphous paracrystalline structures from native crystalline cellulose: A molecular dynamics protocol. Fluid Phase Equilibria 2019;491:56–76. 10.1016/J.FLUID.2019.03.011

7. Poletto M et al. Thermal decomposition of wood: Influence of wood components and cellulose crystallite size. Bioresource Technology 2012;109:148–153. 10.1016/J.BIORTECH.2011.11.122

8. Cui S et al. Exploration of the chemical linkages between lignin and cellulose in poplar wood with 13C and Deuterium dual isotope tracer. Industrial Crops and Products 2022;187:115452. 10.1016/J.INDCROP.2022.115452

9. Štursová M et al. Cellulose utilization in forest litter and soil: identification of bacterial and fungal decomposers. FEMS Microbiology Ecology 2012;80:735–746. 10.1111/j.1574-6941.2012.01343.x

10. Cheshire MV. Origins and stability of soil polysaccharide. Journal of Soil Science 1977;28:1–10. 10.1111/J.1365-2389.1977.TB02290.X

11. Mizuta K, Taguchi S, Sato S. Soil aggregate formation and stability induced by starch and cellulose. Soil Biology and Biochemistry 2015;87:90–96. 10.1016/J.SOILBIO.2015.04.011

12. Ayuso-Fernández I et al. Peroxidase evolution in white-rot fungi follows wood lignin evolution in plants. Proceedings of the National Academy of Sciences of the United States of America 2019;116:17900–17905. 10.1073/PNAS.1905040116

13. Koechli C et al. Assessing fungal contributions to cellulose degradation in soil by using high-throughput stable isotope probing. Soil Biology and Biochemistry 2019;130:150–158. 10.1016/J.SOILBIO.2018.12.013

14. Eaton RA, Hale MDC. Wood: decay, pests and protection., 1st ed. Wood: decay, pests and protection. . London: Chapman and Hall Ltd, 1993.

15. Blanchette RA. Degradation of the lignocellulose complex in wood. Canadian Journal of Botany 1995;73:999–1010. 10.1139/B95-350

16. Cardoso WS et al. Multi-enzyme complex of white rot fungi in saccharification of lignocellulosic material. Brazilian Journal of Microbiology 2018;49:879–884. 10.1016/J.BJM.2018.05.006

17. Okal EJ et al. Mini review: Advances in understanding regulation of cellulase enzyme in white-rot basidiomycetes. Microbial Pathogenesis 2020;147:104410. 10.1016/J.MICPATH.2020.104410

18. Teeri TT. Crystalline cellulose degradation: new insight into the function of cellobiohydrolases. Trends in Biotechnology 1997;15:160–167. 10.1016/S0167-7799(97)01032-9

19. Villares A et al. Lytic polysaccharide monooxygenases disrupt the cellulose fibers structure. Scientific Reports 2017;7:1–9. 10.1038/srep40262

20. Floudas D et al. X-Ray scattering reveals two mechanisms of cellulose microfibril degradation by filamentous fungi. Applied and Environmental Microbiology 2022;88. 10.1128/AEM.00995-22

21. Martinez D et al. Genome, transcriptome, and secretome analysis of wood decay fungus *Postia placenta* supports unique mechanisms of lignocellulose conversion. Proceedings of the National Academy of Sciences of the United States of America 2009;106:1954–1959. 10.1073/PNAS.0809575106

22. Eastwood DC et al. The plant cell wall-decomposing machinery underlies the functional diversity of forest fungi. Science 2011;333:762–765. 10.1126/SCIENCE.1205411

23. Floudas D et al. The paleozoic origin of enzymatic lignin decomposition reconstructed from 31 fungal genomes. Science 2012;336:1715–1719. 10.1126/SCIENCE.1221748

24. Monrroy M et al. Structural change in wood by brown rot fungi and effect on enzymatic hydrolysis. Enzyme and microbial technology 2011;49:472–477. 10.1016/J.ENZMICTEC.2011.08.004

25. Qi J et al. Fungal selectivity and biodegradation effects by white and brown rot fungi for wood biomass pretreatment. Polymers 2023;15:1957. 10.3390/POLYM15081957

26. Hibbett DS, Donoghue MJ. Analysis of character correlations among wood decay mechanisms, mating systems, and substrate ranges in Homobasidiomycetes. Systematic Biology 2001;50:215–242. 10.1080/10635150121079

27. Floudas D. Evolution of lignin decomposition systems in fungi. Advances in Botanical Research 2021;99:37–76. 10.1016/BS.ABR.2021.05.003

28. Floudas D et al. Uncovering the hidden diversity of litter-decomposition mechanisms in mushroom-forming fungi. ISME Journal 2020;14:2046–2059. 10.1038/S41396-020-0667-6

29. Johansson M. A comparison between the cellulolytic activity of white and brown rot fungi. I. The activity on insoluble cellulose. Physiologia Plantarum 1966;19:709–722. 10.1111/J.1399-3054.1966.TB07056.X

30. Highley TL. Influence of carbon source on cellulase activity of white-rot and brown-rot fungi. Wood and Fiber Science 1973;5:50–58.

31. Filley TR et al. Lignin demethylation and polysaccharide decomposition in spruce sapwood degraded by brown rot fungi. Organic Geochemistry 2002;33:111–124. 10.1016/S0146-6380(01)00144-9

32. Liers C et al. Patterns of lignin degradation and oxidative enzyme secretion by different wood- and litter-colonizing basidiomycetes and ascomycetes grown on beech-wood. FEMS microbiology ecology 2011;78:91–102. 10.1111/J.1574-6941.2011.01144.X

33. Yelle DJ et al. Multidimensional NMR analysis reveals truncated lignin structures in wood decayed by the brown rot basidiomycete *Postia placenta*. Environmental microbiology 2011;13:1091–1100. 10.1111/J.1462-2920.2010.02417.X

34. Eichlerová I, Baldrian P. Ligninolytic enzyme production and decolorization capacity of synthetic dyes by saprotrophic white rot, brown rot, and litter decomposing basidiomycetes. Journal of fungi 2020;6:1–22. 10.3390/JOF6040301

35. Csarman F et al. Functional expression and characterization of two laccases from the brown rot *Fomitopsis pinicola*. Enzyme and microbial technology 2021;148. 10.1016/J.ENZMICTEC.2021.109801

36. Suzuki MR et al. Fungal hydroquinones contribute to brown rot of wood. Environmental microbiology 2006;8:2214–2223. 10.1111/J.1462-2920.2006.01160.X

37. Shah F, Mali T, Lundell TK. Polyporales brown rot species *Fomitopsis pinicola*: Enzyme activity profiles, oxalic acid production, and Fe3+-reducing metabolite secretion. Applied and environmental microbiology 2018;84. 10.1128/AEM.02662-17

38. Castaño JD et al. Metabolomics highlights different life history strategies of white and brown rot wood-degrading fungi. mSphere 2022;7. 10.1128/MSPHERE.00545-22

39. Jensen KA et al. An NADH:quinone oxidoreductase active during biodegradation by the brown-rot basidiomycete *Gloeophyllum trabeum*. Applied and environmental microbiology 2002;68:2699–2703. 10.1128/AEM.68.6.2699-2703.2002

40. Arantes V et al. Effect of pH and oxalic acid on the reduction of Fe3+ by a biomimetic chelator and on Fe3+ desorption/adsorption onto wood: Implications for brown-rot decay. International Biodeterioration and Biodegradation 2009;63:478–483. 10.1016/J.IBIOD.2009.01.004

41. Arantes V et al. Lignocellulosic polysaccharides and lignin degradation by wood decay fungi: The relevance of nonenzymatic Fenton-based reactions. Journal of Industrial Microbiology and Biotechnology 2011;38:541–555. 10.1007/S10295-010-0798-2

42. Arantes V, Goodell B. Current understanding of brown-rot fungal biodegradation mechanisms: A review. ACS Symposium Series 2014;1158:3–21. 10.1021/BK-2014-1158.CH001

43. Chirat C, Lachenal D. Effect of hydroxyl radicals on cellulose and pulp and their occurrence during ozone bleaching. Holzforschung 1997;51:147–154. 10.1515/HFSG.1997.51.2.147

44. Tsague FL et al. Study of oxidation of cellulose by Fenton-type reactions using alkali metal salts as swelling agents. Cellulose 2024;31:6643–6661. 10.1007/S10570-024-05970-1

45. Highley TL, Kirk TK, Ibach R. Effect of brown-rot fungi on cellulose. Biodeterioration Research 2. Springer, 1989, 511–525.

46. Espejo E, Agosin E. Production and degradation of oxalic acid by brown rot fungi. Applied and Environmental Microbiology 1991;57:1980. 10.1128/AEM.57.7.1980-1986.1991

47. Kleman-Leyer K et al. Changes in molecular size distribution of cellulose during attack by white rot and brown rot fungi. Applied and environmental microbiology 1992;58:1266–1270. 10.1128/AEM.58.4.1266-1270.1992

48. Li GY et al. Chemical compositions, infrared spectroscopy, and X-ray diffractometry study on brown-rotted woods. Carbohydrate Polymers 2011;85:560–564. 10.1016/J.CARBPOL.2011.03.014

49. Hastrup ACS et al. Differences in crystalline cellulose modification due to degradation by brown and white rot fungi. Fungal Biology 2012;116:1052–1063. 10.1016/J.FUNBIO.2012.07.009

50. Zhu Y et al. Non-enzymatic modification of the crystalline structure and chemistry of Masson pine in brown-rot decay. Carbohydrate Polymers 2022;286:119242. 10.1016/J.CARBPOL.2022.119242

51. Fackler K et al. Localisation and characterisation of incipient brown-rot decay within spruce wood cell walls using FT-IR imaging microscopy. Enzyme and Microbial Technology 2010;47:257–267. 10.1016/J.ENZMICTEC.2010.07.009

52. Agarwal UP. 1064 nm FT-Raman spectroscopy for investigations of plant cell walls and other biomass materials. Frontiers in Plant Science 2014;**5**:103972. 10.3389/FPLS.2014.00490

53. Agarwal UP. Analysis of cellulose and lignocellulose materials by Raman spectroscopy: A review of the current status. Molecules 2019;24:1659. 10.3390/MOLECULES24091659

54. Agarwal UP. Beyond crystallinity: Using Raman spectroscopic methods to further define aggregated/supramolecular structure of cellulose. Frontiers in Energy Research 2022;10:857621. 10.3389/FENRG.2022.85762

55. Wiley JH, Atalla RH. Band assignments in the Raman spectra of celluloses. Carbohydrate Research 1987;160:113–129. 10.1016/0008-6215(87)80306-3

56. Agarwal UP et al. Probing crystallinity of never-dried wood cellulose with Raman spectroscopy. Cellulose 2016;23:125–144. 10.1007/S10570-015-0788-7

57. Agarwal UP et al. New cellulose crystallinity estimation method that differentiates between organized and crystalline phases. Carbohydrate Polymers 2018;190:262–270. 10.1016/J.CARBPOL.2018.03.003

58. Zhang X, Li L, Xu F. Polarized Raman spectroscopy for determining the orientation of cellulose microfibrils in wood cell wall. Cellulose 2023;30:75–85. 10.1007/S10570-022-04915-W

59. Pence I, Mahadevan-Jansen A. Clinical instrumentation and applications of Raman spectroscopy. Chemical Society reviews 2016;45:1958. 10.1039/C5CS00581G

60. Cordero E et al. In-vivo Raman spectroscopy: from basics to applications. Journal of biomedical optics 2018;23:1. 10.1117/1.JBO.23.7.071210

61. Lu L et al. Resonance Raman scattering of β-carotene solution excited by visible laser beams into second singlet state. Journal of Photochemistry and Photobiology B: Biology 2018;179:18–22. 10.1016/J.JPHOTOBIOL.2017.12.022

62. Jiao C, Xiong J. Accessibility and morphology of cellulose fibres treated with sodium hydroxide. BioResources 2014;9:6504–6513.

63. Isogai A, Saito T, Fukuzumi H. TEMPO-oxidized cellulose nanofibers. Nanoscale 2011;3:71–85. 10.1039/C0NR00583E

64. Tang Z et al. TEMPO-oxidized cellulose with high degree of oxidation. Polymers 2017;9. 10.3390/POLYM9090421

65. Heinze T. Cellulose Chemistry and Properties: Fibers, Nanocelluloses and Advanced Materials, 1st ed. Springer Cham, 2015.

66. Oh SY et al. Crystalline structure analysis of cellulose treated with sodium hydroxide and carbon dioxide by means of X-ray diffraction and FTIR spectroscopy. Carbohydrate Research 2005;340:2376–2391. 10.1016/J.CARRES.2005.08.007

67. TJ W. Amplification and direct sequencing of fungal ribosomal RNA genes for phylogenetics. PCR protocols : A guide to methods and applications 1990;315–322.

68. Gardes M, Bruns TD. ITS primers with enhanced specificity for basidiomycetes - application to the identification of mycorrhizae and rusts. Molecular Ecology 1993;2:113–118. 10.1111/J.1365-294X.1993.TB00005.X

69. Saitou N, Nei M. The neighbor-joining method: a new method for reconstructing phylogenetic trees. Molecular Biology and Evolution 1987;4:406–425. 10.1093/OXFORDJOURNALS.MOLBEV.A040454

70. Katoh K, Rozewicki J, Yamada KD. MAFFT online service: multiple sequence alignment, interactive sequence choice and visualization. Briefings in bioinformatics 2019;20:1160–1166. 10.1093/BIB/BBX108

71. Waterhouse AM et al. Jalview Version 2—a multiple sequence alignment editor and analysis workbench. Bioinformatics 2009;25:1189–1191. 10.1093/BIOINFORMATICS/BTP033

72. Kumar S et al. MEGA X: Molecular evolutionary genetics analysis across computing platforms. Molecular biology and evolution 2018;35:1547–1549. 10.1093/MOLBEV/MSY096

73. Toplak M et al. Quasar: Easy machine learning for biospectroscopy. Cells 2021;10. 10.3390/CELLS10092300

74. Troein C et al. OCTAVVS: A graphical toolbox for high-throughput preprocessing and analysis of vibrational spectroscopy imaging data. Methods Protoc 2020;3. 10.3390/mps3020034

75. O’haver T. An introduction to signal processing in chemical measurement. Article in Journal of Chemical Education 1991;68. 10.1021/ed068pA147

76. R. Core Team. R: A language and environment for statistical computing. R Foundation for Statistical Computing . 2021. R Foundation for Statistical Computing, 2021.

77. Rousseeuw PJ. Silhouettes: A graphical aid to the interpretation and validation of cluster analysis. Journal of Computational and Applied Mathematics 1987;20:53–65. 10.1016/0377-0427(87)90125-7

78. Agarwal UP, Reiner RS, Ralph SA. Cellulose I crystallinity determination using FT-Raman spectroscopy : univariate and multivariate methods. Cellulose 2010;17:721– 733. 10.1007/S10570-010-9420-Z

79. Agarwal UP et al. Contributions of crystalline and noncrystalline cellulose can occur in the same spectral regions: Evidence based on Raman and IR and its implication for crystallinity measurements. Biomacromolecules 2021;22:1357–1373. 10.1021/ACS.BIOMAC.0C01389

80. Bates D et al. Fitting linear Mixed-effects models using lme4. Journal of Statistical Software 2015;67:1–48. 10.18637/JSS.V067.I01

81. Shapiro SS, Wilk MB. An analysis of variance test for Normality (complete samples). Biometrika 1965;52:591. 10.2307/2333709

82. Friendly M. HE Plots for Multivariate Linear Models. Journal of Computational and Graphical Statistics 2007;16:421–444. 10.1198/106186007X208407

83. Larsson E, Larsson KH. Phylogenetic relationships of russuloid basidiomycetes with emphasis on aphyllophoralean taxa. Mycologia 2003;95:1037–1065. 10.1080/15572536.2004.11833020

84. Howell C et al. Effects of hot water extraction and fungal decay on wood crystalline cellulose structure. Cellulose 2011;18:1179–1190. 10.1007/S10570-011-9569-0

85. Witomski P, Olek W, Bonarski JT. Changes in strength of Scots pine wood (*Pinus silvestris* L.) decayed by brown rot (*Coniophora puteana*) and white rot (*Trametes versicolor*). Construction and Building Materials 2016;102:162–166. 10.1016/J.CONBUILDMAT.2015.10.109

86. Moreau C et al. Lytic polysaccharide monooxygenases (LPMOs) facilitate cellulose nanofibrils production. Biotechnology for Biofuels 2019;12:1–13. 10.1186/S13068-019-1501-0

87. Bissaro B, Eijsink VGH. Lytic polysaccharide monooxygenases: enzymes for controlled and site-specific Fenton-like chemistry. Essays in Biochemistry 2023;67:575–584. 10.1042/EBC20220250

88. Munzone A et al. Expanding the catalytic landscape of metalloenzymes with lytic polysaccharide monooxygenases. Nature Reviews Chemistry 2024;8:106–119. 10.1038/s41570-023-00565-z

89. Boulos S, Nyström L. UPLC-MS/MS investigation of β-glucan oligosaccharide oxidation. Analyst 2016;141:6533–6548. 10.1039/C6AN01125J

90. Qin RC et al. Preparation of cellulose nanofibers from corn stalks by Fenton reaction: A new insight into the mechanism by an experimental and theoretical study. Journal of Agricultural and Food Chemistry 2023;71:1907–1920. 10.1021/ACS.JAFC.2C08475

91. Howell C et al. Temporal changes in wood crystalline cellulose during degradation by brown rot fungi. International Biodeterioration & Biodegradation 2009;63:414–419. 10.1016/J.IBIOD.2008.11.009

92. Schwarze FWMR, Engels J, Mattheck C. Fungal strategies of wood decay in trees, 1st ed. Springer Berlin Heidelberg, 2000.

93. Nakasone KK. Cultural studies and identification of wood-inhabiting Corticiaceae and selected Hymenomycetes from North America. Schweizerbart Science Publishers, 1990.

94. Ginns JH, Lefebvre MNL. Lignicolous corticioid fungi (Basidiomycota) of North America : systematics, distribution, and ecology. Brittonia 1995;47:92. 10.2307/2807250

95. Otjen L, Blanchette RA. *Xylobolus frustulatus* decay of oak: Patterns of selective delignification and subsequent cellulose removal. Applied and environmental microbiology 1984;47:670–676. 10.1128/AEM.47.4.670-676.1984

96. Riley R et al. Extensive sampling of basidiomycete genomes demonstrates inadequacy of the white-rot/brown-rot paradigm for wood decay fungi. Proceedings of the National Academy of Sciences of the United States of America 2014;111:9923–9928. 10.1073/PNAS.1400592111

97. Zhang X et al. Obtaining pure spectra of hemicellulose and cellulose from poplar cell wall Raman imaging data. Cellulose 2017;24:4671–4682. 10.1007/S10570-017-1486-4

98. Bock P et al. Infrared and Raman spectra of lignin substructures: Dibenzodioxocin. Journal of Raman Spectroscopy 2020;51:422–431. 10.1002/JRS.5808

99. Gao W et al. Reliable and realistic models for lignin content determination in poplar wood based on FT-Raman spectroscopy. Industrial Crops and Products 2022;182:114884. 10.1016/J.INDCROP.2022.114884

